# The mechanosensitive Piezo1 orchestrating angiogenesis is essential in bone fracture repair

**DOI:** 10.1101/2019.12.23.887661

**Authors:** Peng Chen, Gangyu Zhang, Shan Jiang, Yile Ning, Bo Deng, Xianmei Pan, Silin Liu, Yu He, Lei Zhang, Rentao Wan, Zhiming Wu, Qi He, Jiang Yin, Haibin Wang, Jing Li

**Affiliations:** The First Affiliated Hospital Guangzhou University of Chinese Medicine, Guangzhou, 510405, China; Lingnan Medical Research Center, Guangzhou University of Chinese Medicine, Guangzhou, 510405, China; The First School of Clinical Medicine, Guangzhou University of Chinese Medicine, Guangzhou, 510405, China; School of Pharmaceutical Sciences, Guangzhou University of Chinese Medicine, Guangzhou, 510405, China; The affiliated Cancer Hospital, Guangzhou Medical University, Guangzhou, China, 510095

**Keywords:** *Piezo1* channels, endothelial knockout mice, angiogenesis, bone fracture, vascular biology

## Abstract

Mechanical ion channel protein *Piezo1* play vital roles in angiogenesis which has been proved to be high importance in varieties of biological processes. Bone formation in the fracture repair requires oxygen and nutrients from new blood vessels generated from fractured lesion. Understanding the underlying mechanisms linking angiogenesis and bone formation must be of great value for improved fracture healing. Here we employed mice with genetically modified endothelial specific depletion of *Piezo1* channels to explore the hypothesis that *Piezo1* is vital to the initiation of fracture healing. In this study, we demonstrated that *Piezo1* expression and wide distribution along the bone and impaired endothelial *Piezo1* channels result in derangements in bone fracture repair. Intriguingly, the calcium activated proteolytic caplain activity severely disrupted during vascularization, precluded osteoblast maturation and mineralization and subsequently the phosphorylated *PI3K-AKT* reduction. Furthermore, *Piezo1* endothelial disruption impaired *Notch* signaling in bone union. These data collectively suggest that *Piezo1* channels serve as a basis for clinical strategies to improve bone regeneration and treat delayed or nonunion in bone fracture.

## Introduction

The skeleton is a complex multifunctional organ system and bone fractures are the traumatic injuries of the most common large-organs in humans. The bone repair following fractures is a postnatal regeneration in skeletal system and usually an efficient and rapid process involving both biological and mechanical aspects which aims at restoring the fractured bone to its biomechanical function and pre-damaged structure. Unfortunately, approximately 5%-10% injured bones manifesting delayed healing and/or nonunion results in restrained movements (1–3). The impaired fracture healing are often associated with a variety of risk factors including nutritional, biological as well as physical factors. Among these, neovascularization is believed to be crucial in normal bone healing(4–11). Several molecular players, for example, vascular endothelial growth factor (VEGF) signaling(12–14), hypoxia(15, 16), matrix metalloproteases (17–19), fibroblast growth factor (FGF) (20, 21) and Notch signaling (22, 23) had been intensively explored. However, the mechanism orchestrating angiogenesis during fracture healing has not been very well defined.

The recently discovered mechanically activated cationic ion channel *Piezo1*, has proved as a Ca^2+^ permeable transmembrane protein activated by physical force (24, 25). It consists of a three blade assembled pore unit and formed a propeller like structure and is involved in multiple biological functions, which determined cells in responding to touch, cell volume, or stretching of the surrounding tissue through converting mechanical force into electrochemical signals (26–33). In previous studies, endogenous *Piezo1* channels are highly expressed in vasculature where it directly sense the physiological shear stress imposed on endothelial cells (ECs) which have the key roles in vascular development (34, 35). Moreover, its function has been verified in physical exercise, vascular remodeling and lymphatic vascular development (36–41). More recently, *Piezo1* channels, as the critical mechanotransducer in controlling mechanosensitivity in osteogenesis and mechanical force dependent bone formation has been confirmed (42, 43).

Given the vital importance of *Piezo1* in vascular endothelial cells, we speculate *Piezo1* might play an essential role in bone formation and remodeling during bone fracture healing. We systematically detected the function of *Piezo1* in this process using conditional *Cre-Lox*-mediated depletion of *Piezo1* in the endothelium (*Piezo1*^*ΔEC*^ mice) and compared with their littermates (*Piezo1*^*fl/fl*^ mice). Our findings revealed that *Piezo1* coupled to *Notch* signaling pathway to determine bone remodeling.

## Results

### Vascular expression and distribution of *Piezo1* in bone tissue

It has been reported that *Piezo1* ion channels are widely expressed in a variety of cells and organs. To examine the vascular arrangement of *Piezo1* channels in bone, we employed knock-in reporter mice wherein a tandem-dimer Tomato (tdT) fluorescent protein is tagged to the C-terminus of the *Piezo1* channels, which has been confirmed that the expression levels and patterns are of the same as the endogenous *Piezo1* channels (35, 44). Immunostaining showed bone vasculature labeled with endomucin (*EMCN*) that is a membrane-bound glycoprotein expressed on the surface of endothelial cells and the *Piezo1*-tdT labelling with an anti-RFP antibody as indicated by the representative confocal images (Fig. 1A). *Piezo1*-tdT was found localized in a large proportion to endomucin-labeled endothelial cells. In line with this result, western blot analysis revealed that the *Piezo1*-tdT fusion protein was readily detected in bone tissue from *Piezo1-*tdT mice and no signal was observed in lysates obtained from *Piezo1* wild-type (*Piezo1*^+/+^) mice (Fig. 1B). The data suggest that endothelial *Piezo1* is abundant in bone vasculature.

**Fig. 1.**
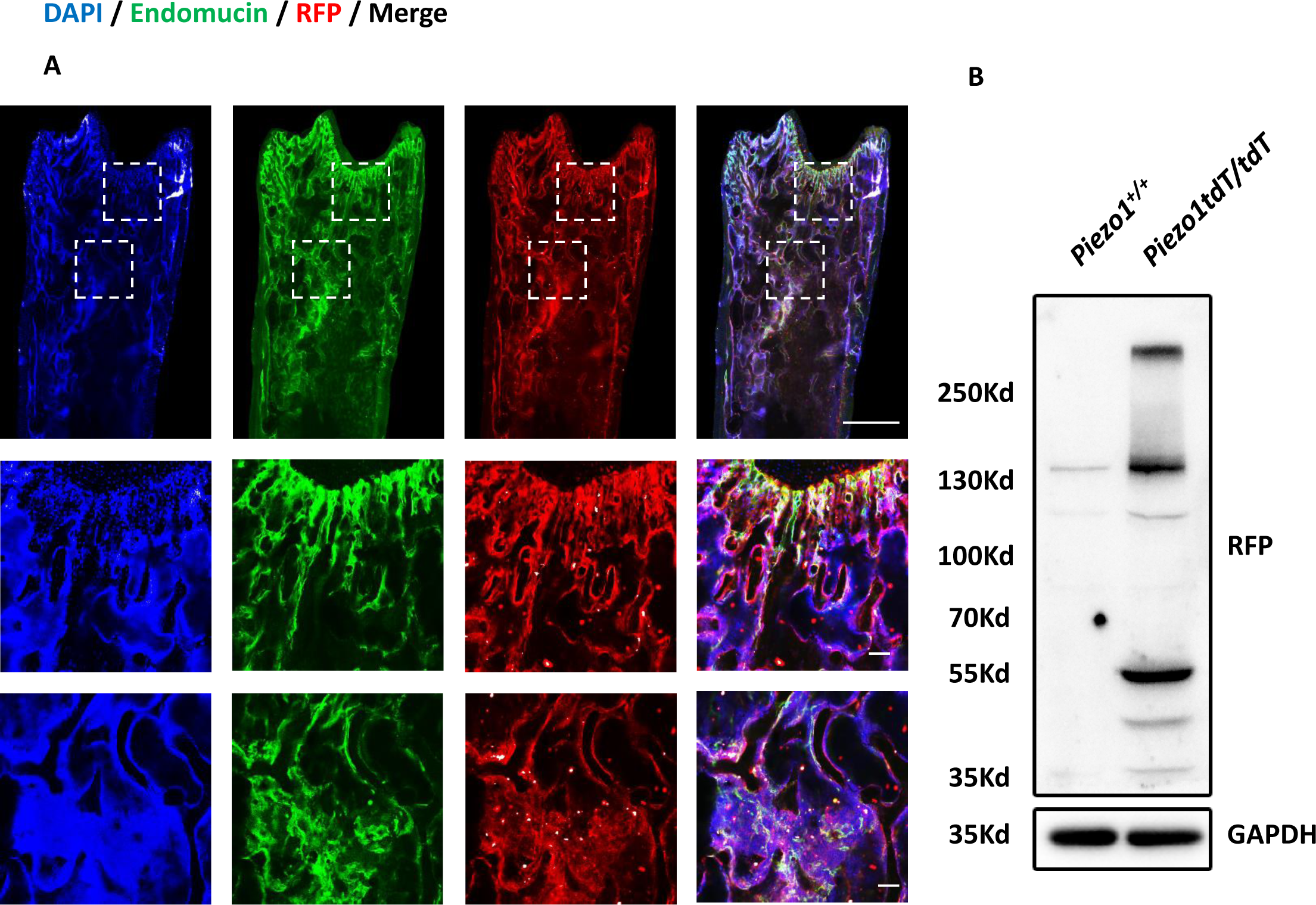
*Piezo1*-tdT expression and distribution in the femora of *Piezo1*-tdT reporter mice. **(A)** Representative confocal images of the femora in *Piezo1-*tdT mice stained with *EMCN* (green), *RFP* (red) and DAPI (blue). Insets were magnified images of the areas enclosed by white dashed boxes. Scale bars, 1mm (top); 100 µm (middle and bottom). (B) Bone tissue lysates, prepared from Piezo1+/+ and Piezo1-tdT mice, were resolved by SDS-PAGE and western blots probed with an antibody against red fluorescent protein (which reacts with the tdTomato moiety). Detection of *GAPDH* was used as loading control (n = 3).

### *Piezo1*^*ΔEC*^ mice generation and validation

To detect the endothelial *Piezo1* specific disruption in mice we freshly isolated liver endothelial cells from *Piezo1*^*ΔEC*^ and *Piezo1*^*fl/fl*^ mice. Gel electrophoresis of PCR end-point products, quantitative RT-PCR analysis and Yoda1 induced calcium entry in liver endothelial cells from *Piezo1*^*ΔEC*^ and *Piezo1*^*fl/fl*^ mice revealed the specific knockout of *Piezo1* in endothelium (Fig. 2A-D). In contrast, ATP induced calcium entry showed no difference in liver endothelial cells from *Piezo1*^*ΔEC*^ and *Piezo1*^*fl/fl*^ mice (Fig. 2E). Thus, we next focus on investigating the function of endothelial *Piezo1* in skeletal system.

**Fig. 2.**
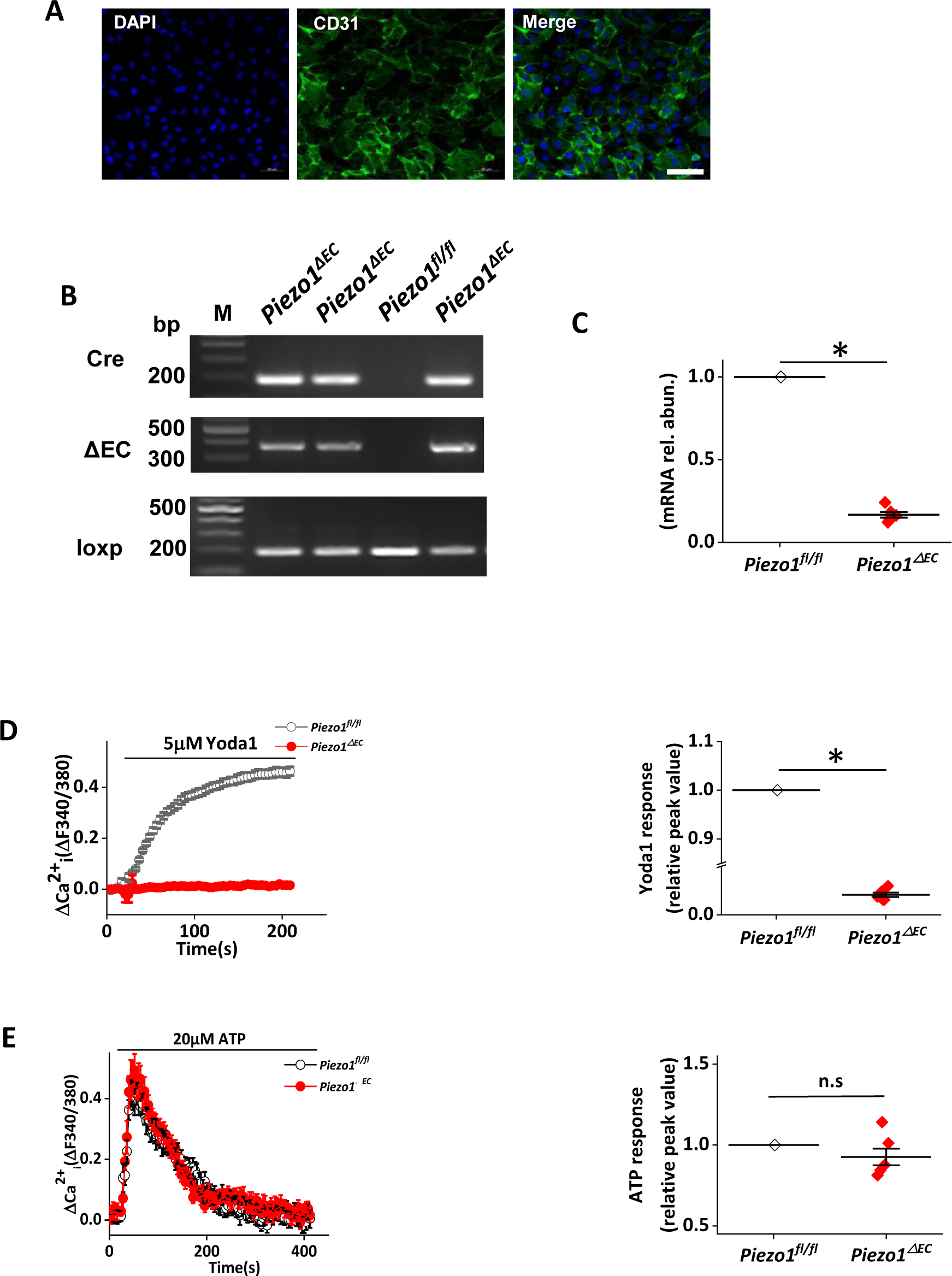
Validation of *Piezo1* endothelial depletion mice (*Piezo1*^*ΔEC*^ mice) Liver endothelial cells which we freshly isolated from *Piezo1*^*ΔEC*^ and *Piezo1*^*fl/fl*^ mice (**A**) Immunofluorescence of liver endothelial cells stained with anti-CD31antibody (green) and DAPI (blue). Scale bar, 20 μm. (**B**) Gel electrophoresis of PCR end-point products. Template was DNA from liver endothelial cells isolated from three *Piezo1*^ΔEC^ mice and one *Piezo1*^*fl/fl*^ mouse. Presence of the cre recombinase transgene was shown in the top row (Cdh5-cre, 191 bps); deletion of *Piezo1* gene was shown in the middle row (ΔEC, 379 bps); presence of LoxP sequence in *Piezo1* gene was shown in the bottom row (*Piezo1*Flox, 189 bps). (**C**) Quantitative RT-PCR analysis of *Piezo1* mRNA abundance in liver endothelial cells from *Piezo1*^*fl/fl*^ and *Piezo1*^*ΔEC*^ mice (n=6). (**D**) On the left, representative traces of change(Δ) in intracellular Ca^2+^ in liver endothelial cells which from *Piezo1*^*ΔEC*^ and *Piezo1*^*fl/fl*^ mice upon application of 5 µM Yoda1. On the right, mean data for the Yoda1 response at 200 s in *Piezo1*^*ΔEC*^ and *Piezo1*^*fl/fl*^ mice of liver endothelial cells (n=6). (**E**).On the left, representative traces of change (Δ) in intracellular Ca^2+^ in *Piezo1*^*ΔEC*^ and *Piezo1*^*fl/fl*^ mice liver endothelial cells upon application of 20 µM ATP (n=6). On the right, mean data for the ATP response at 80 s in *Piezo1*^*ΔEC*^ and *Piezo1*^*fl/fl*^ mice of liver endothelial cells (n=6).

### *Piezo1*^*ΔEC*^ mice show severely impaired bone formation

To determine the potential role of *Piezo1* during bone fracture repair, we performed stabilized femur fractures on *Piezo1*^ΔEC^ and their littermates (*Piezo1*^*fl/fl*^) when mice were injected with tamoxifen one week later. As shown in macro-images (Fig. 3A), callus has been well-formed and remodeling at the fracture ends at 21 days post fracture (DPF) and 42 DPF in *Piezo1*^*fl/fl*^ mice. In contrast, callus was still defective at the fracture site by 21 DPF and callus resorption has not finished even by 42 DPF. Similarly, X-ray and 3D μCT in *Piezo1*^*fl/fl*^ fractures illustrated external callus formed completely at the fracture site by 21 DPF, followed by the completion of bone remodeling by 42 DPF, suggesting bone fracture well unified. Even though *Piezo1*^*ΔEC*^ mice demonstrated remarkable callus formation along the periosteum extending away from the fracture line, no bridging callus was clearly observed by 21 DPF. And even more, radiographic evident up to 42 DPF still highlighted a clear radiolucent gap between broken ends (Fig. 3B and 3C). Further analysis of the periosteal callus around the site of fracture revealed more bony trabecular and foci of mineralization in axial μCT at 21 and 42 DPF in *Piezo1*^*fl/fl*^ mice than i *Piezo1*^*ΔEC*^ mice (Fig. 3D). Consistently, coronal view from 3D μCT data of *Piezo1*^*fl/fl*^ fractures showed a nearly complete bridging of bony calluses by 21 DPF, followed by complete bridging by 42 DPF. *Piezo1*^*ΔEC*^ fractures presented with a large gap between fracture cortices sties up to and beyond 42 DPF (Fig. 3E). Quantitatively, at 21 and 42 DPF in *Piezo1*^*ΔEC*^ mice, trabecular bone volume fraction (BV/TV), trabecular thickness (Tb.Th), trabecular number (Tb.N) were significantly decreased while trabecular separation (Tb.Sp) increased compared to their littermates which indicated the fracture union delayed or fracture nonunion. The data suggest the obvious nonunion developed in bone fracture in *Piezo1* endothelial depletion mice.

**Fig. 3.**
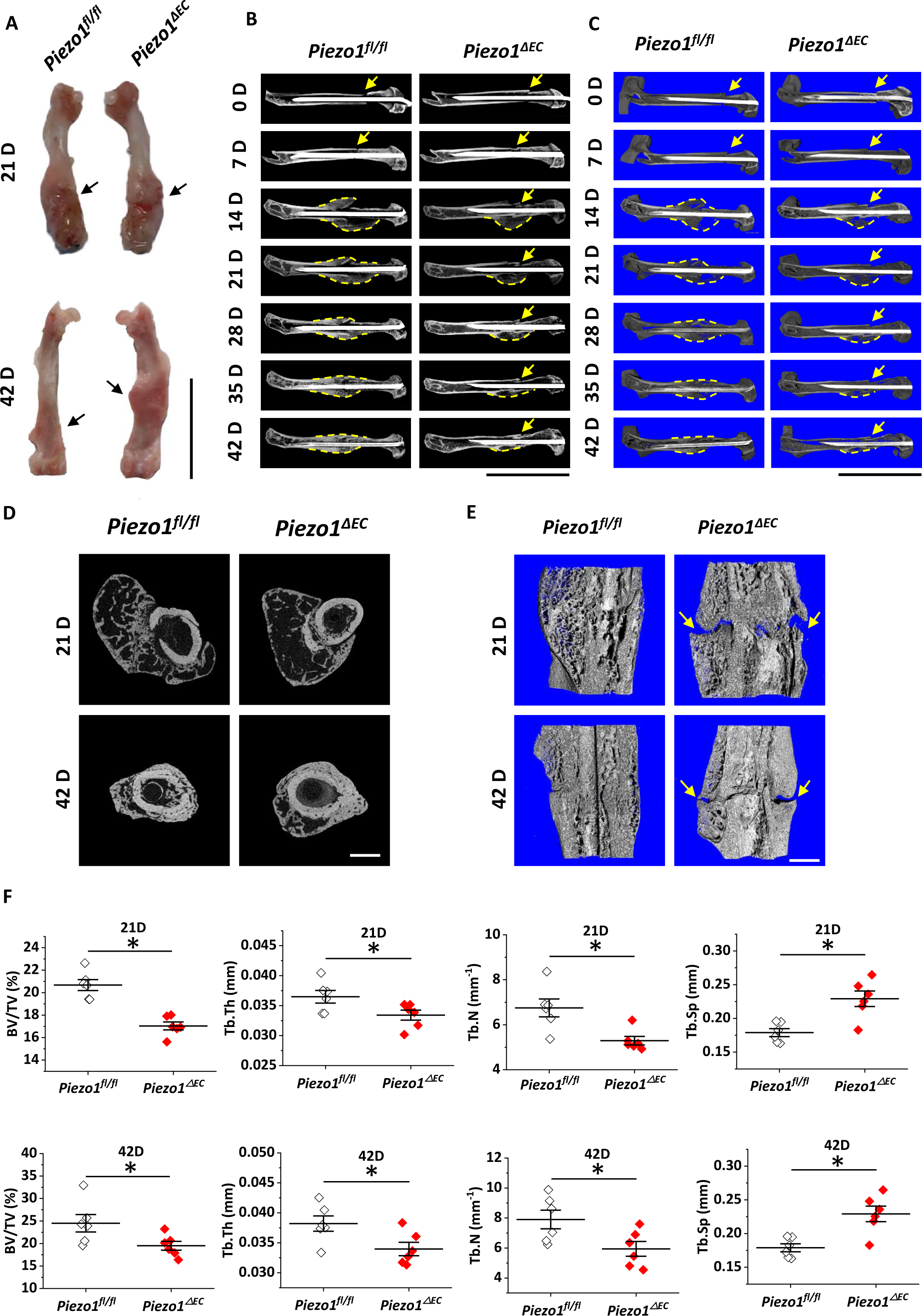
*Piezo1*^*ΔEC*^ mice show severely impaired bone formation. **(A)** Representative macroscopic photograph of fractured femora showed callus has been well-formed and well remolded in *Piezo1*^*fl/fl*^ mice at 21DPF, 42DPF compared to *Piezo1*^*ΔEC*^ mice (black arrows denote fracture callus). Scale bars: 1 cm. n = 6 mice per genotype per time point. **(B, C)** A real-time X-ray and 3D μCT radiographic comparison of 2 representative femoral fractures from *Piezo1*^*fl/fl*^ and *Piezo1*^*ΔEC*^ mice at 0, 7, 14, 21, 28, 35 and 42 DPF suggested persistent fracture lines (yellow arrows) at 42 DPF, revealing an established fracture nonunion in *Piezo1*^*ΔEC*^. Scale bars: 1 cm. n = 6 mice per genotype per time point. **(D)** Representative μCT axial images of fractured femurs at 21 and 42 DPF in *Piezo1*^*fl/fl*^ mice showed bony trabecular and foci of mineralization compared to *Piezo1*^*ΔEC*^ mice. Scale bars, 1 mm. n = 6 for each genotype. **(E)** Representative coronal view from 3D μCT data at 21 and 42 DPF in *Piezo1*^*fl/fl*^ mice demonstrated that well-formed and well-remodeling bony callus while apparent radiolucent space (yellow arrows) between broken cortices in *Piezo1*^*ΔEC*^ mice. Scale bars: 1 mm. n = 6 for each genotype. **(F)** BV/TV, Tb.N, and Tb.Th were significantly higher in *Piezo1*^*fl/fl*^ mice at 14 and 42 DPF while Tb.Sp was lower. \**P < 0.05* by 2-tailed, unpaired Student’s test. Results are expressed as mean±SEM.

### *Piezo1*^*ΔEC*^ mice show less expression of endomucin (*EMCN*), *RUNX2 and* Osterix (*Osx*) during bone formation

Given that endothelial *Piezo1* channels are crucial in vascular formation in development, we hypothesized that depletion of *Piezo1* in endothelial cells might affect angiogenesis and subsequent osteogenesis in bone repair. We performed immunostaining on regional sections of fractured femurs. As demonstrated in Fig. 4A and B, *EMCN*, a sialoprotein expressed by non-arterial endothelial cells, was less abundant around fracture site in *Piezo1*^*ΔEC*^ mice than that in *Piezo1*^*fl/fl*^ mice, indicating impaired new blood vessels formation. To further determine whether osteogenesis factors, *RUNX2*, which has a crucial role in the early determination stage of the osteoblast lineage and *Osterix* (*Osx*), which regulates the late stage of osteoblast differentiation and bone formation were affected by endothelial *Piezo1* knockout mice, we found both *RUNX2* and *OSX* were dramatically decreased in *Piezo1*^*ΔEC*^ mice compared with *Piezo1*^*fl/fl*^ mice, as shown in Fig. 4C and Fig. 4E and quantified the fluorescent intensity in Fig. 4D and Fig. 4F. The data suggest that endothelial *Piezo1* has key roles in soft and hard callus formation through angiogenesis which providing the blood and nutrients in bone fracture healing process.

**Fig. 4.**
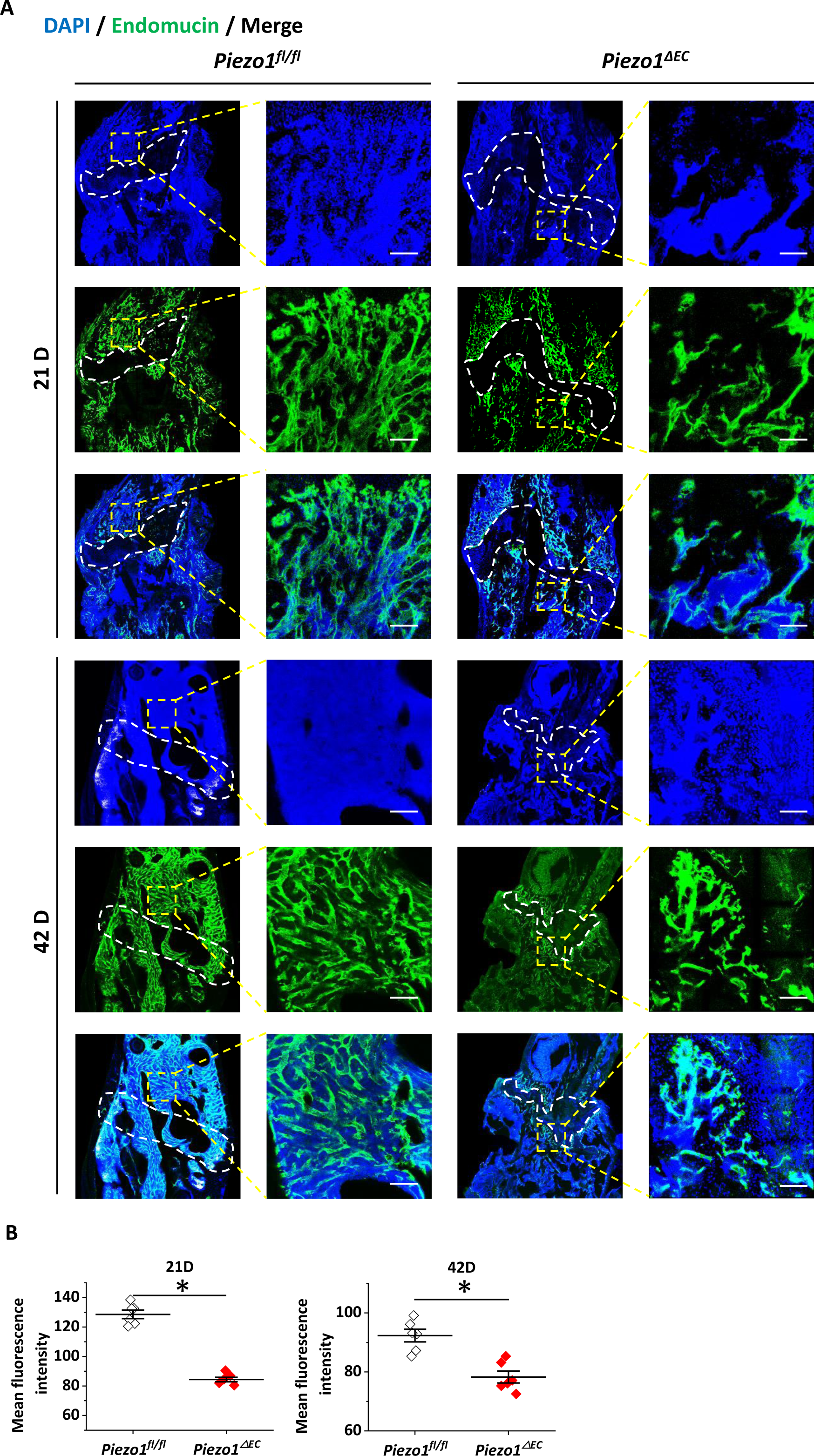
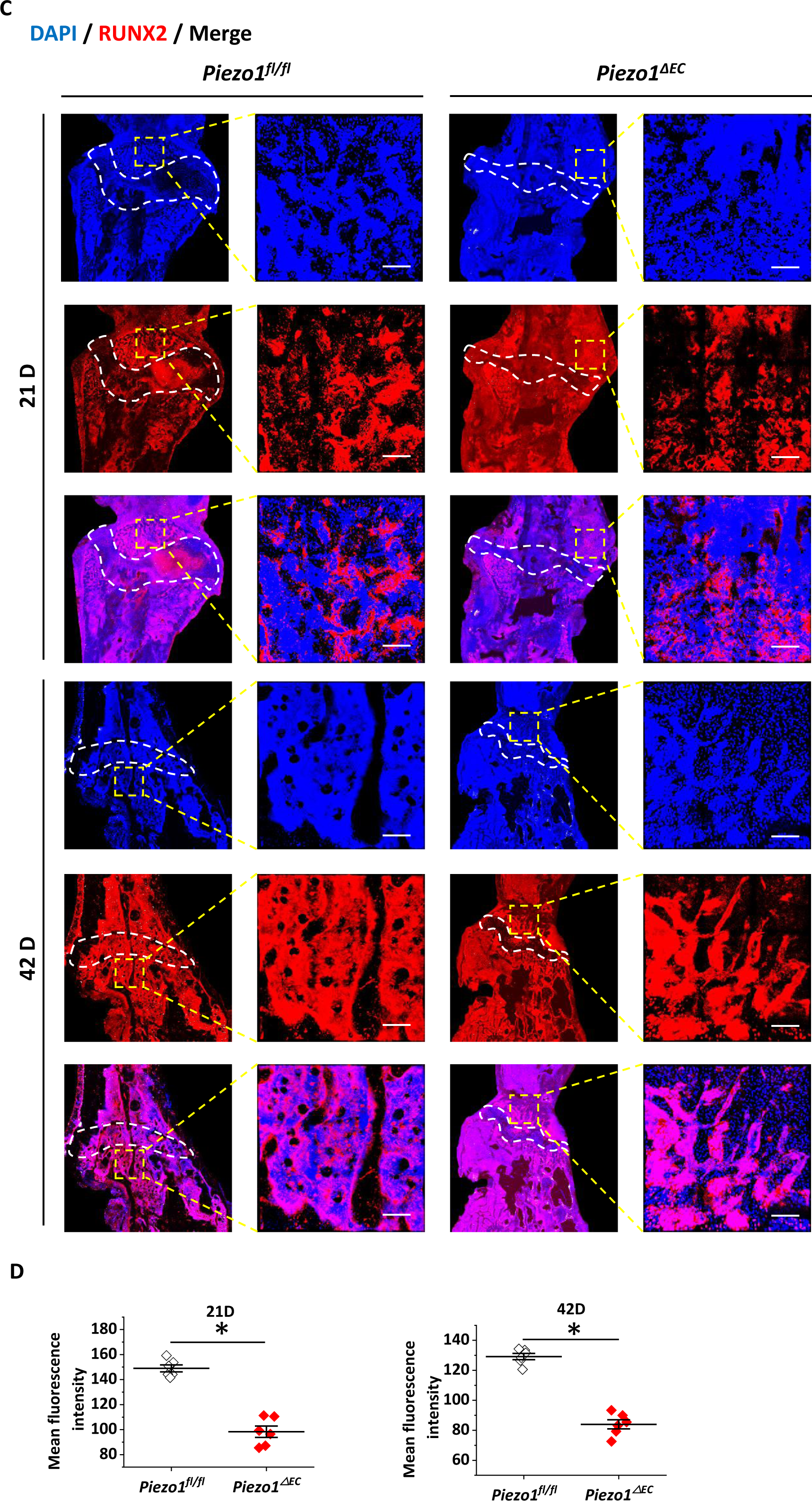
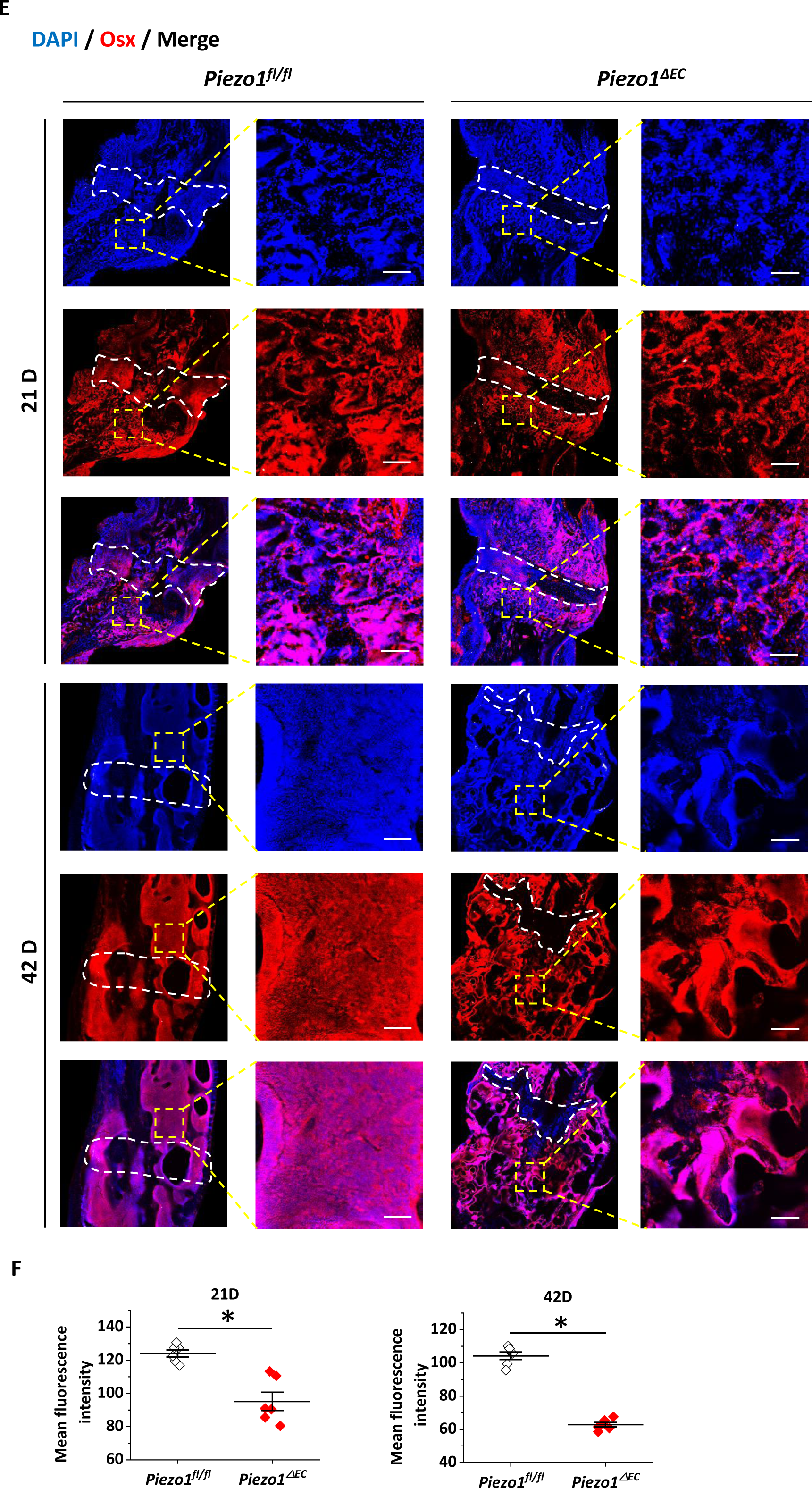
*Piezo1*^*ΔEC*^ mice show less expression of *EMCN*, *RUNX2*, and *Osx* during bone formation. **(A, B)** Representative confocal images and quantification of *Piezo1*^*fl/fl*^ and *Piezo1*^*ΔEC*^ mice femora stained with *EMCN* (green) and DAPI (blue). Insets were magnified images of the fractured areas enclosed by white dashed boxes. Scale bars, 100 µm. n = 6 mice per genotype per time point. **(C, D)** Representative confocal images and quantification of *Piezo1*^*fl/fl*^ and *Piezo1*^*ΔEC*^ mice femora stained with *RUNX2* (red) and DAPI (blue). Insets were magnified images of the fractured areas enclosed by white dashed boxes. Scale bars, 100 µm. n = 6 mice per genotype per time point. **(E, F)** Representative confocal images and quantification of *Piezo1*^*fl/fl*^ and *Piezo1*^*ΔEC*^ mice femora stained with *Osx* (red) and DAPI (blue). Insets were magnified images of the fractured areas enclosed by white dashed boxes. Scale bars, 100 µm. n = 6 mice per genotype per time point. \**P < 0.05* by 2-tailed, unpaired Student’s test. Results are expressed as mean±SEM.

### Loss of endothelial *Piezo1* impairs bone fracture healing by histological examination

To better examine the role of *Piezo1* during bone fracture repair, histological staining was employed to determine the calluses formed between the fracture gap and periosteum. At 21 DPF, H&E staining exhibited complete continuity of the cortex with mature lamellar structure in *Piezo1*^*fl/fl*^ mice (Fig. 5A upper). Meanwhile, ABH/OG and SFO/FG staining revealed that the morphology and the sizes of cartilage-like or bony-like calluses formed between the fracture gap and periosteum. Activated osteocytes (bright blue with pericellular rings in Fig. 5B upper) and cartilage-containing areas (red area in Fig. 5C upper) also reflected the strong osteoblastic activity, whereas no healing evident in *Piezo1*^*ΔEC*^ mice. In place of normal healing, *Piezo1*^*ΔEC*^ mice osteotomies remained a clear broken gap at broken sites and no obvious osteocytes and ossification centers. At 42 DPF, bony calluses were reduced to normal cortical bone by bone remodeling process in *Piezo1*^*fl/fl*^ mice. However, fracture line was still evident even the occurrence of adipose vesicles in *Piezo1*^*ΔEC*^ mice. (Fig. 5A-C lower parts).

**Fig. 5.**
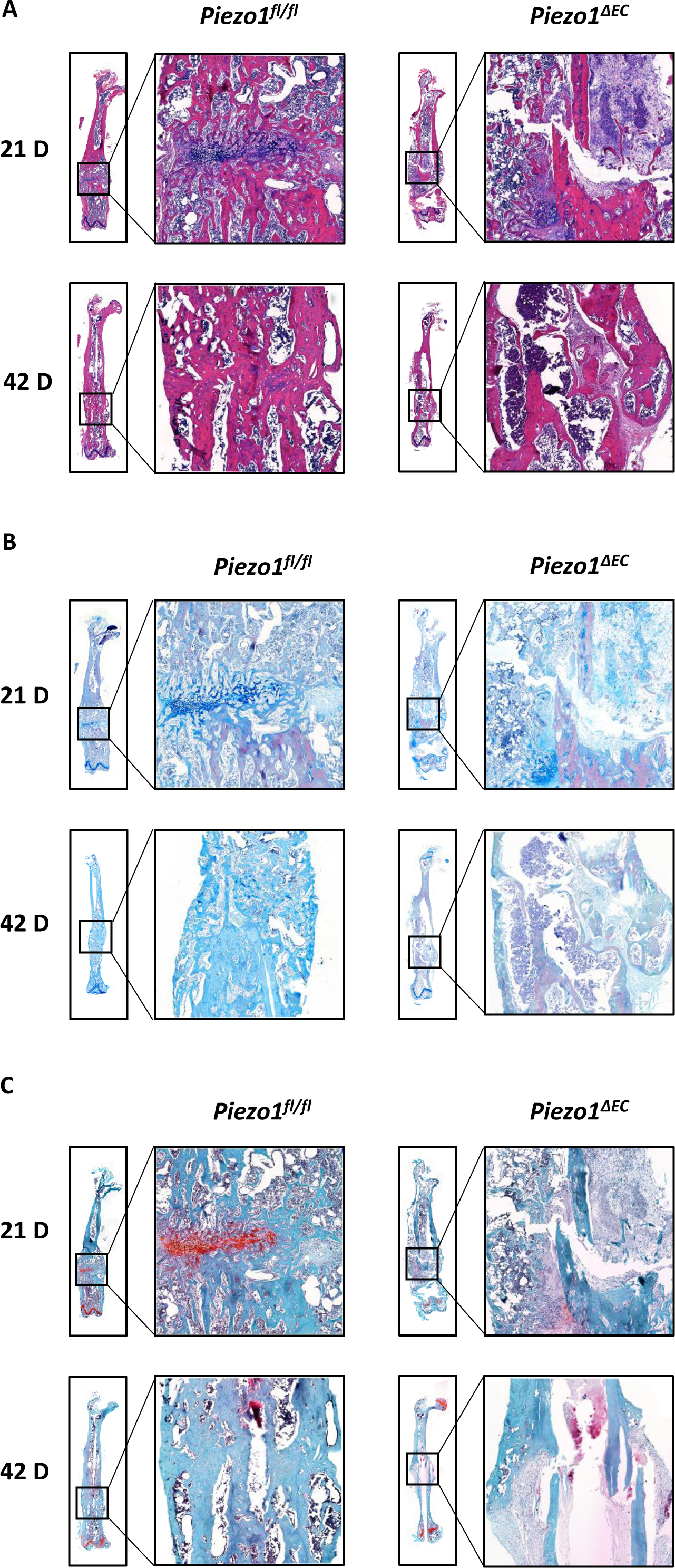
Loss of endothelial *Piezo1* results in histological changes. **(A)** Representative images of H&E staining of femora at 21 and 42 DPF between *Piezo1*^*fl/fl*^ and *Piezo1*^*ΔEC*^ mice. n = 6 mice per genotype per time point. **(B)** Representative images of ABH/OG staining of femora at 21 and 42 DPF between *Piezo1*^*fl/fl*^ and *Piezo1*^*ΔEC*^ mice. n = 6 mice per genotype per time point. **(C)** Representative images of SFO/FG staining of femora at 21 and 42 DPF between *Piezo1*^*fl/fl*^ and *Piezo1*^*ΔEC*^ mice. n = 6 mice per genotype per time point.

### Loss of *Piezo1* severely impairs angiogenesis

Blood supply to the fracture sites is essential for bone fracture repair. In order to determine the role of *Piezo1* on angiogenesis in the process of bone fracture healing, we employed Microfil perfusion and found that visible new blood vessels were formed around the fracture site in *Piezo1*^*fl/fl*^ mice at 21 DPF compared with *Piezo1*^*ΔEC*^ mice. Furthermore, the blood vessel turned out to be close to normal in *Piezo1*^*fl/fl*^ mice at 42 DPF while the blood vessel in *Piezo1*^*ΔEC*^ mice remained interrupted (Fig. 6A). Quantitatively, not only the vessel volume (fraction), but also the vessel surface were significantly lower in *Piezo1*^*ΔEC*^ mice than in their littermates both by 21 and 42 DPF (Fig. 6B). In accordance with our expectations, these data reveal that *Piezo1* is able to orchestrate angiogenesis during fracture healing.

**Fig. 6.**
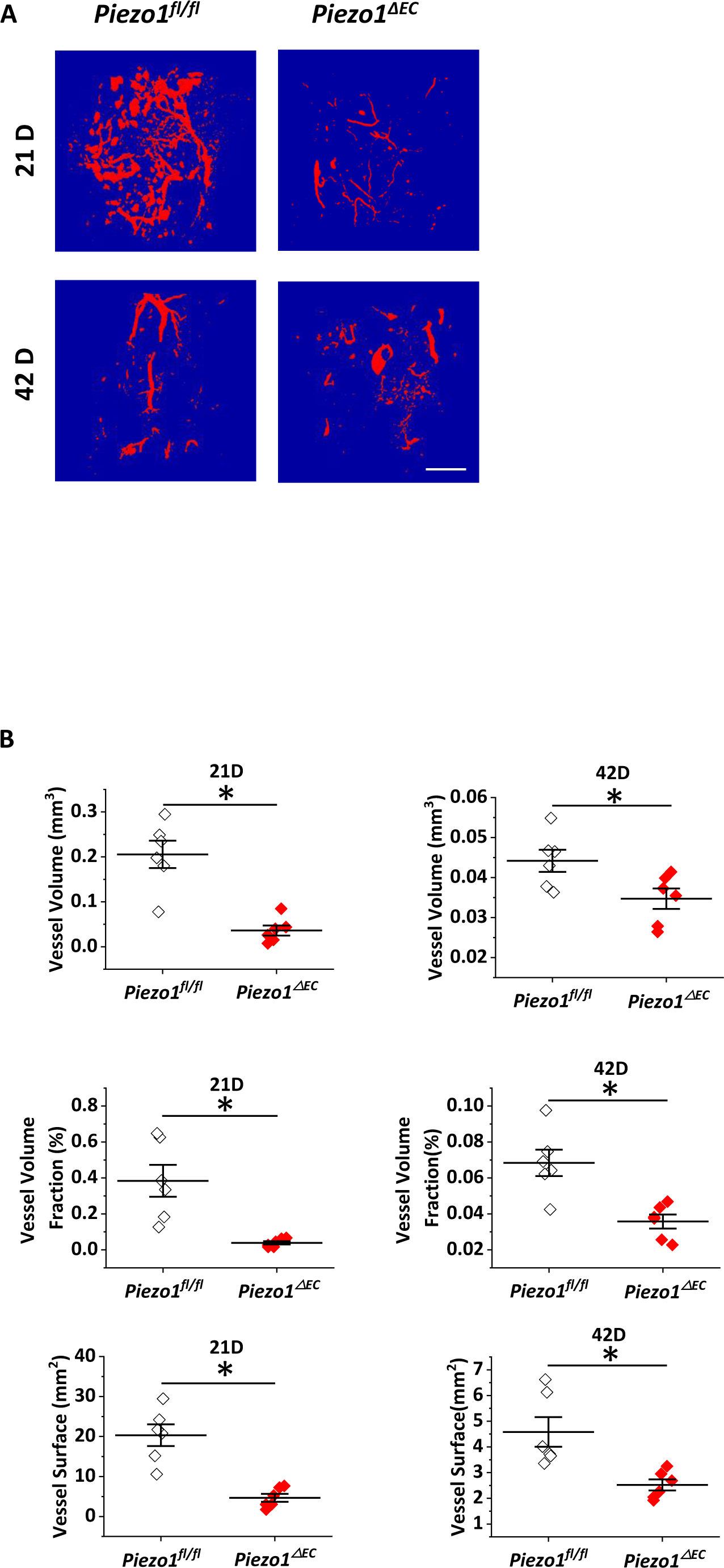
Loss of endothelial *Piezo1* severely impairs angiogenesis. Representative µCT based angiographic images **(A)** and quantification of vessel volume, vessel volume fraction and vessel surface area **(B)**. Scale bar, 1 mm. n = 6 per group per time point. \**P < 0.05* by 2-tailed, unpaired Student’s test. Results are expressed as mean±SEM.

### Loss of *Piezo1* remarkably reduces biomechanical properties

We next sought to determine whether the altered fracture repair processes have the impact on bone structure. To this end, we analyzed the mechanical properties of fracture femora by three-bending test. As expected, we found yield load and bending stiffness were significantly decreased in samples of 21 DPF in *Piezo1*^*ΔEC*^ mice compared to *Piezo1*^*fl/fl*^ mice. Although the two parameters were slightly enhanced by 42 DPF in *Piezo1*^*ΔEC*^ mice compared to 21 DPF, there were still remarkably lower than *Piezo1*^*fl/fl*^ mice (Fig. 7). These data indicate that biomechanical properties are indeed reduced with the loss of endothelial *Piezo1*.

**Fig. 7.**
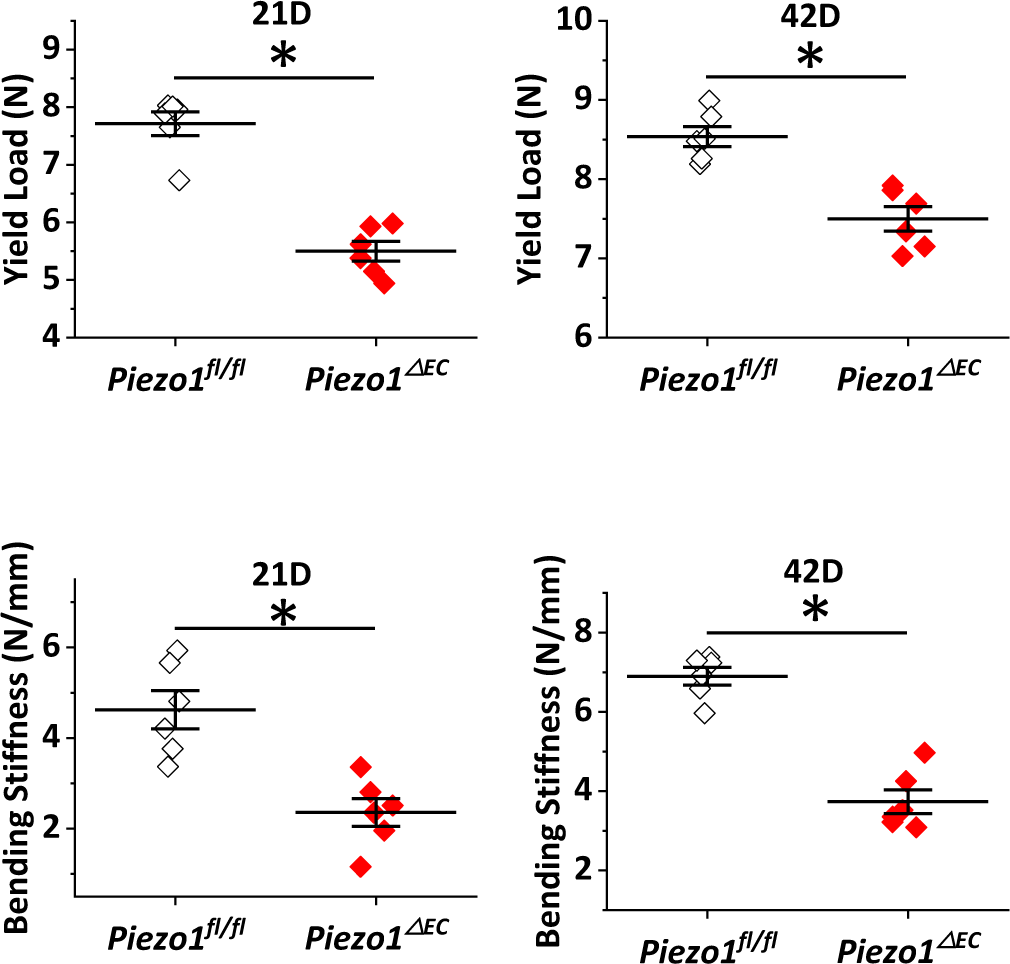
Loss of endothelial *Piezo1* remarkably reduces biomechanical properties. Yield load and bending stiffness were significantly decreased in samples of 21 DPF and 42 DPF in *Piezo1*^*ΔEC*^ mice compared to *Piezo1*^*fl/fl*^ mice. n = 6 mice per genotype per time point. \**P < 0.05* by 2-tailed, unpaired Student’s test. Results are expressed as mean ±SEM.

### *Piezo1* endothelial depletion impairs calpain activity

To gain insight into the mechanistic understanding of the role of *Piezo1* channels in fracture healing, we next performed the calpain activity assay, given that *Piezo1* channels can activate calpain, a cytoplasmic cysteine protease which requires calcium ions for its function and subsequently mediates a variety of intracellular processes including cleaving focal adhesion proteins and cytoskeletal substrates to impair angiogenesis (34). We found that there was significantly decreased calpain activity in *Piezo1*^*ΔEC*^ mice compared to *Piezo1*^*fl/fl*^ mice, suggesting that *Piezo1* activation of calpain was responsible for bone reunion (Fig. 8A).

**Fig. 8.**
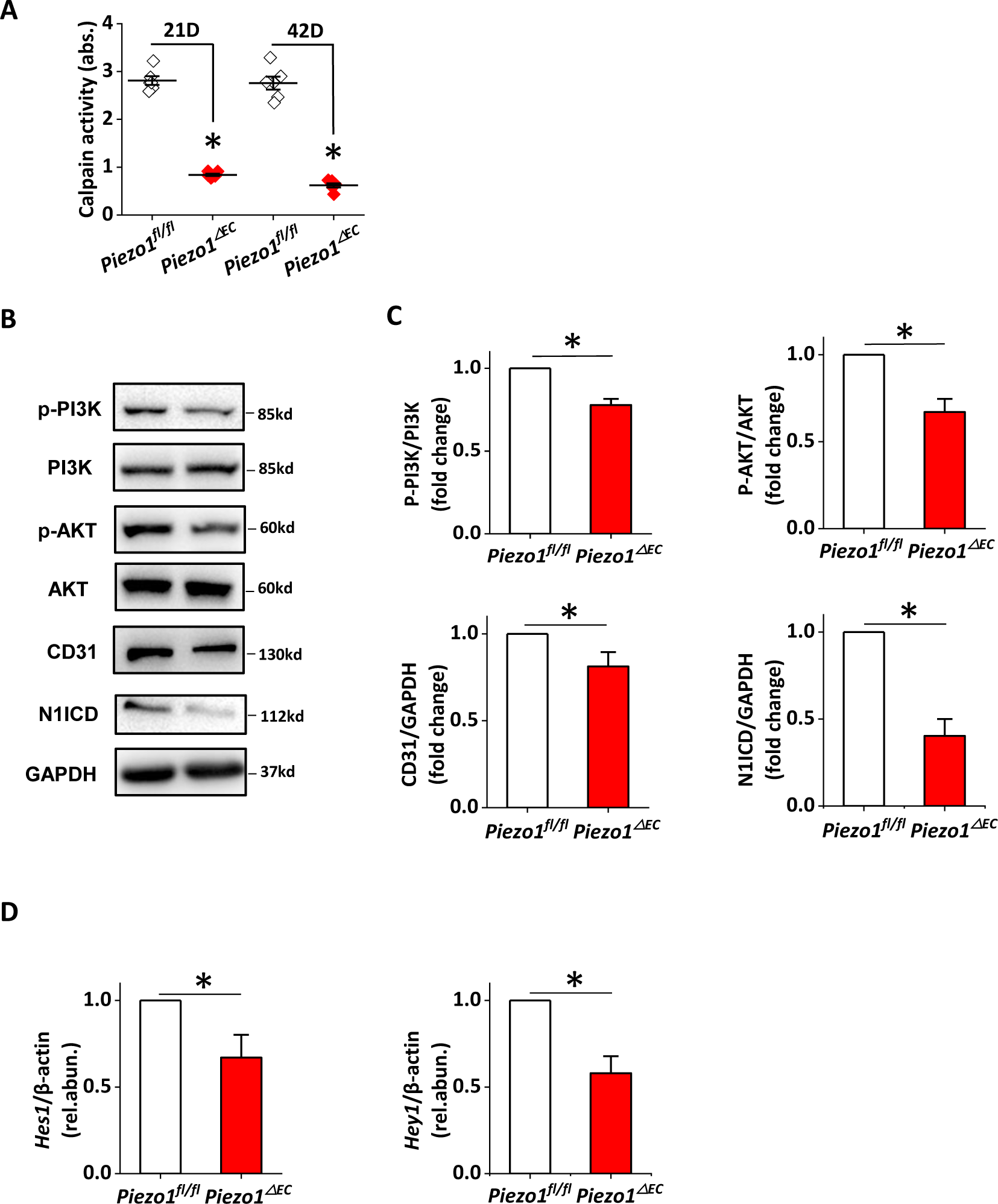
Loss of endothelial *Piezo1* reduces calpain activity and impairs Notch signaling pathway. **(A)** Calpain activity measurements of tissues from bone fracture lesions at day 21 and 42, respectively in *Piezo1*^*ΔEC*^ mice compared to *Piezo1*^*fl/fl*^ mice. n = 6 mice per genotype per time point. **(B-F)** Protein levels, determined by western blotting, and mean data for *p-PI3K*, *p-AKT*, *CD31* and *N1ICD* in *Piezo1*^*ΔEC*^ mice compared to *Piezo1*^*fl/fl*^ mice. n = 3 each. **(G-H)** mRNA levels of *Hes1* and *Hey1* were detected by real-time PCR in *Piezo1*^*ΔEC*^ mice compared to *Piezo1*^*fl/fl*^ mice. n = 6 each.

### *Piezo1* endothelial depletion couples to *PI3k-AKT* pathway and *Notch* signaling in fracture repair

*Piezo1* serves as a non-selective cationic ion channel that allows Ca^2+^ entry and initiates a Ca^2+^ dependent downstream signals. Previous studies have demonstrated that Ca^2+^ entry mechanism could lead to phosphorylation of *PI3K-AKT* and thus regulate proliferation, migration and angiogenesis in endothelial cells (45, 46) and promote osteoblast differentiation and mineralization(47). A mechanical stimulation mediated *Piezo1*-*Akt* pathway in osteocytes is required for bone homeostasis (48) In line with these reports, we identified that the phosphorylation of *PI3K* and *AKT* was dramatically decreased in *Piezo1*^*ΔEC*^ mice compared to *Piezo1*^*fl/fl*^ mice at the fracture lesions. Furthermore, *CD31*(*PCAM-1*), and *Notch1* intracellular domain (*N1ICD*), the key players in promoting flow induced angiogenesis, were downregulated in *Piezo1*^*ΔEC*^ mice compared to *Piezo1*^*fl/fl*^ mice (Fig. 8B) and quantified in (Fig. 8C). Consistently, the expression levels of *Hes1* and *Hey1*, the *Notch1* regulated genes, showed dramatic reduction in *Piezo1*^*ΔEC*^ mice compared with in *Piezo1*^*fl/fl*^ mice (Fig. 8D). Taken together, the data suggest that endothelial deficiency of mechanosensitive *Piezo1* channels significantly impairs the Ca^2+^ dependent bone angiogenesis and ossification in bone fractural healing processes.

## Discussion

To investigate whether *Piezo1* ion channels, which contribute crucially to vascularization in early development in mice, are playing roles in bone fracture healing process we examined mice with stabilized fracture in endothelial specific deletion of *Piezo1* channels. These studies revealed that the exact *Piezo1* expression architecture of the bone vasculature and demonstrated strikingly that *Piezo1* is an important player in vascularization in bone fracture reunion and that the impaired Ca^2+^ entry mediated proteolysis calpain activation upon *Piezo1* endothelial depletion. Furthermore, we found that a positive feedback relationship between *Piezo1* channel and *PI3K-AKT* downstream pathway as well as *CD31* and *Notch* signaling in regulating bone new vessel formation and bone ossification.

It has been widely reported that bone fracture healing contains a more complex process and is different from a wound repair which creates a fibrous scar in a soft tissue. The steps of fracture repair include hematoma formation where broken of blood vessels result in inflammatory cells release and coagulation cascade activation and soft and hard callus formation where intramembranous ossification, the internal callus becomes mineralized when angiogenesis occurred and finally the fractured bone remodeling by returning vascular structure and replace the damaged bone(49). In the past years, extensive studies have led to the discovery of a large number of growth factors that stimulate angiogenesis, which are very important in bone development and normal fracture healing. *Notch* signaling is one of the most well investigated angiogenic factors in the mammalian skeletal system and has been proved that manipulation of *Notch* receptors lead to desired successful fracture repair in femoral fracture and cortical bone defects in mice. Consistently our experiments showed that depletion of *Piezo1* reduced *Notch1* expression and further proved *Notch* signaling pathways are indispensable for vascularization, callus formation and mineralization while the bone was injured. These findings provide also the notion that *Piezo1* combined with *Notch* is the key mechanism where angiogenesis and osteogenesis are tightly coupled in bone repair.

There are increasing recognition of calpain mediated signaling pathway in angiogenesis. Several seminal studies have revealed that *VEGF* induced calpain dependent activation of *PI3K-AKT* cascade, subsequent release of nitric oxide and generation of physiological angiogenesis. We have reported that when HUVECs overexpressing *Piezo1*-WT-GFP plasmid-were applied with flow accumulation of *Piezo1*-WT-GFP at the leading apical lamellipodia were achieved which facilitating HUVECs to migrate and thus form the new blood vessels. The harmonization of cytoskeleton and focal adhesion was controlled importantly by calpain. We further reported that at embryonic E9.5 and 10.5 stages global disruption of *Piezo1* led to the decreased calpain activity. In line with this findings it is a new paradigm that the important roles of *Piezo1* in angiogenesis and bone repair and remodeling during fracture.

In present studies we focused only on the endothelial function of *Piezo1* in bone fracture healing, As mentioned previously, the critical roles of *Piezo1* channels in bone development have been published by two groups (42, 43) and based on the widely expression of *Piezo1* in red blood cells, macrophages, monocyte and leucocytes, we postulated that the role of *Piezo1* channels in impaired fracture repair may be functionally involved a more complex picture. Although there are many unanswered questions we are still facing the great opportunity to explore the precise role of bone EC subsets and blood vessels in physiological and pathophysiological environments, which will improve our mechanistic understanding of the skeletal system from the cellular and molecular aspects. Further extensive studies are needed to investigate the crosstalk between different signaling regulations and to elucidate the potential therapeutic applications aimed at preventing bone loss or stimulating fracture healing.

In summary, our results indicate that *Piezo1* channels play an important role in regulating Ca^2+^ entry mechanism and local vascular growth and provide valuable niche signals for osteogenesis and in reducing *Notch1* expression in fracture healing. Although mouse models of fracture healing very closely reproduce the process found in humans, they may not be exactly the same. As yet pharmacologically intervention of *Piezo1* channels has limited but with high resolution cryo-EM structure identification of *Piezo1* channels there will be more effective small molecules to be synthesized and clinically applied to the patients. Nevertheless, our data provide insight into how mechanical sensitive endothelial cells can promote bone fracture healing processes, particularly in regard to vascularization through a *Piezo1* regulated mechanism.

## Materials and Methods

### Genetically modified mice

All animal studies were carried out with approval from Animal Care and Use Committee of Guangzhou University of Chinese Medicine. The *Cre/loxP* system was introduced to obtain the conditional knockout mice. Endothelial-cell-specific gene deletions mice were created and characterized by intercrossing Cdh5 (PAC)-CreERT2 transgenic mice (a gift from Ralf Adams, University of Münster, Münster, Germany) with conditional mutants containing loxP flanked *Piezo1* (*Piezo1*^*lox/lox*^). In order to achieve Cre activity and gene deletion, offspring mice of 8-week old were treated with 500 mg tamoxifen (Sigma, T5648) intraperitoneally (i.p.) for 5 consecutive days. Although experimenters were not blinded when dividing mice into experimental groups, we definitely conduct fracture performance and end-point harvesting under blindness. Mice with the fused sequence for tdTomato (*Piezo1-*tdT) at the C-terminus of *Piezo1* and *Piezo1*^*lox/lox*^ were purchased from The Jackson Laboratory. PCR test were used to genotype all mice. Protocols for PCR experiment and sequences for primers will be available upon request.

### Establishment of open femoral shaft transverse fracture mice model

A unilateral (right side) open transverse femur fracture was performed according to previously described protocols with modifications (50). Briefly, mice were anesthetized with 1% pentobarbital (50 mg/kg) by intraperitoneal injection. We depilated the right leg with an electric shaver and cleaning the local area with an alcohol wipe, prior to making an incision above the right anterolateral femur. Muscle were separated by blunt dissection and exposed the middle shaft femur. When patella was flipped outwardly, a 27 gauge syringe needle was inserted into the medullary cavity paralleling with the long axis of the femur through the center of trochlear groove. The needle was then pulled out, and transverse diaphyseal fracture was created in the middle of the femur with fine scissors. Then, a 26 gauge syringe needle was aseptically introduced into the bone marrow cavity through the fracture line to achieve the fracture stabilization. The tail of needle was trimmed carefully to protect patellofemoral joint movement. The periosteum remained intact and soft tissues were well protected. Before closing the wound, neither partial fracture nor comminuted fracture was confirmed. A 4-0 silk suture was used to stitch the wound. After recovery from anesthesia, mice were placed back to their home cages with free movement. In the first three days after operation, penicillin ointment was used to avoid wound infection while buprenorphine was administered in drinking water for pain relief. Fracture repair was followed radiographically using a micro CT (Skyscan 1176 model, Bruker) weekly under anesthesia.

### High-resolution 3D confocal imaging acquisition and quantitative analysis

The injured femora were dissected at 21 or 42 days after operation. During the procedure, all the surrounding muscles attached to the femur were carefully excoriated to avoid causing any mechanical damage to bones and callus tissue. The intramedullary needles were also removed carefully. We then strictly followed the standard protocol for immunofluorescence (51). Fresh bone samples were immediately fixed in ice-cold 4% (wt/vol) paraformaldehyde (PFA) solution for 4h in a 15 mL conical tube sitting on the ice. We washed the bone tissues in 3 changes of sterile 1X PBS (7 mL) for 5 min at 4 °C each time with persistent agitation to make sure that all of the PFA was completely removed. Then, we added ice-cold 0.5 M EDTA (10 mL, PH 7.4) and incubated the mixture at 4 °C under constant shaking up to 24h (this decalcification step could be assessed by pressing the sample with a forceps), followed by bone samples being washed 3 times with 7 mL of PBS and each time for 5 min at 4 °C under constant agitation. PBS was removed completely from the tube and 10 mL of ice-cold cryoprotectant (CPT) solution was added for 24h. After completely remove CPT, bone samples were transferred to a tissue mold and immediately embedded with freshly made embedding solution (EBM) of approximately 5 mL without any bubbles or solid particles (the tissue mold must be filled up completely with EBM). The samples were incubated for half an hour at RT to ensure EBM totally solidification followed by 45 min of incubation at 60°C in water bath. Subsequently, we transferred the samples to - 80°C for storage overnight, before placed the samples for section with a thickness of 50 µm (the cryotome was set at low speed of 20-40%).

For immunofluorescence staining, we rehydrated the sections by incubating bone slides with 200 µL of PBS for 5 min at RT and then 200 µL 0.3% (vol/vol) Triton X-100 permeabilization (PRM) solution at RT for 20 min. Then, we used donkey serum (5%) to block sections for half an hour at RT and incubated them overnight at 4 °C with 200 µL of primary antibody solutions: *RFP* (600-401-379, 1:100; Rockland), *Endomucin* (*EMCN*, sc-65495, 1:100; Santa Cruz), *RUNX2* (ab23981, 1:100; Abcam), *Osterix* (*OSX*, ab22552, 1:100; Abcam). After primary antibody incubation, slides were rinsed 3 changes with PBS for 5 min each and incubated with 200 µL of fluorescent-conjugated secondary antibody at RT for 60 min while light avoidance. After that, sections were washed with 3 changes of 1X PBS for 5 min each. Then, the nuclei were counterstained with freshly prepared 4’, 6-diamidino-2-phenylindole (DAPI). We preformed all immunofluorescence experiments at least three times. We used Leica TCS SPEII confocal microscope for imaging samples with tile z-stack sequential scanning. We quantified all high resolution confocal images with ImageJ software.

### High-resolution *in-vivo* µCT of *Piezo1*^*fl/fl*^ and *Piezo1*^*ΔEC*^ mice

The operated femora were imaged weekly from the first day after surgery to 42 days under anesthesia with high-resolution *in-vivo* µCT scanner (Skyscan 1176, Bruker, MA, USA). The scanner was set up with an AI (0.5 mm) filter. The tube voltage was set to 50 kV with the beam current 500 μA. The scanning angular rotation was set as 180° while the angular increment was 0.50° with isotropic voxel size of 9.02 μm. We employed image reconstruction software (NRecon V1.6.9), data analysis software (CTAn v1.9) and three-dimensional model visualization software (µCTVol v2.0) for analyzing the parameters of the fracture site. Two-dimensional morphometric analysis of the cortical bone and three-dimensional histomorphometric evaluation of the trabecular bone were performed with established cross-sectional images of the femora. Trabecular bone region of interest (ROI) was drawn commencing at the upper 500 slices to the lower 500 slices at the center of fracture line. Respectively, we segmented the images into trabecular bone and bone marrow. The parameters of trabecular bone volume fraction (BV/TV), trabecular thickness (Tb.Th), trabecular number (Tb.N) and trabecular separation (Tb.Sp) were further analyzed.

### Histological analysis

Femora samples were harvested from *Piezo1* mice at 21 DPF and 42 DPF, respectively. Under the condition of euthanasia, fracture femora were dissected and then fixed in 4% PFA. We decalcified the tissues with 0.5M EDTA (pH 7.4) for 2 weeks and dehydrated them in an ascending ethanol series, then in xylene for clear. Tissues were processed to be embedded in paraffin and 6 µm longitudinally thickness sections were cut by microtomes. All fixation and decalcification steps were carried out at RT.

For Hematoxylin & Eosin (H&E) stain, slides were deparaffinized and rehydrated by distilled water. Then, we placed slides into Hematoxylin solution for 5-10 min and washed them in tap water for 1 to 5 min until sections became blue. Sequentially, slides were soaked in Eosin solution for about 30 seconds and then rinsed in tap water when water became clear again. Then, slides were dehydrated by 2 times of 95% ethanol, cleared in 2 times of xylene for 5 min before mounting on coverslips using 3-4 drops of Permount.

For Alcian Blue Hematoxylin/ Orange G (ABH/OG) stain, slides were deparaffinized and rehydrated by distilled water. Then, slides were immersed into acid-alcohol for 30 seconds and drain briefly and stained with Alcian Blue Hematoxylin working solution for 40 min. Sections were next rinsed gently with tap water until excess stain stops leaching from tissue. Then, slides were differentiated in acid-alcohol for 3 second and washed by distilled water 2 times. Sequentially, slides were dipped into 95% ethanol for 1 minute and placed in Eosin solution for 90 seconds. Then, slides were mounted on coverslips with Permount following by dehydrated by 3 changes of 95% ethanol and 2 changes of 100% ethanol and cleared in 3 changes of xylene.

For Safranin Orange/Fast Green (SFO/FG) stain, slides were baked at 60°C overnight and cooled down at RT about 20-30 min. Then, we deparaffinized the slides and hydrated them with 70% ethanol. Following that, we stained the slides by immersion in freshly prepared working Weigert’s hematoxylin for 7 min and quickly rinsed them with running tap water until it runs clear (approximately 10 min). Then, slides were immersed in working 0.08% fast green solution for 3 min and washed immediately by 1% acetic acid for 10 seconds. Slides were then stained with 0.1% safranin O solution for 5 min. Ultimately, slides were dehydrated by passage through 2 times of 95% ethanol and 100% ethanol for 3 min each and then cleared in 2 times of xylene for 5 min. Slides were then coverslipped by Permount.

### CT based microangiography

In order to characterize the blood vessels during bone fracture healing process at 21 DPF and 42 DPF respectively, CT based angiography of microfil perfusion was introduced in this study. Briefly, experimental animals were anesthetized with 1% pentobarbital (50 mg/kg) by intraperitoneal injection and positioned supine. After exposing thoracic cavity, we irrigated the vascular system through 0.9% saline solution containing heparin sodium (100 U/mL) by inserting a blunt 22 gauge needle into the left ventricle. Immediately, we cut the right atrium open to ensure systemic blood outflows and continued flushing when the blood was thoroughly expelled from circulation. Microfil MV–122 (Flow Tech, USA), a low-viscosity radiopaque polymer, was used to identify the circulatory system followed by the specimens were fixed with 10% neutral buffered formalin under constant pressure. In order to confirm entirely filled with the vasculature, we assessed the extravascular pooling by judging the appearance of both the liver and the last organ to be perfused prior to extravasation through the right atrium. Mice were included in our study only with the complete hepatic blanching prior to Microfil obtained, or the contrast was very clear, or without extravascular pooling occurrence. Mice were kept at 4 °C and maintained overnights to enable contrast agent polymerization. Femora were then dissected and immersed in 10% neutral buffered formalin for 4 days for totally fixation. Following that, all fracture samples were decalcified for 48h in a 0.5 M EDTA (PH 7.4) and then acquired images again by μCT imaging system (Skyscan 1172). CTAn was used to reconstruct and quantify the vascular model.

### Biomechanical testing

3-point bending test was introduced to determine the bone mechanical properties with Electroforce 3220 (Bose Corporation, USA). Experiment was conducted with the load point in displacement control, moving at a speed of 0.03 mm per second. Each sample was put in the same orientation with the cranial surface sitting on the supporting surface. Yield load (N) and bending stiffness (N-mm^2^) were analyzed in accordance with the force and displacement data acquired in the tests.

### Calpain activity assay

Calpain activity assay kit (Abcam, ab65308) was used to measure calpain activity with detection of cleavage of calpain substrate. Briefly, mice bone samples were dissected, homogenized and re-suspended in the provided extraction buffer. Protein concentration was determined and the inputs were standardized according to total protein content. Subsequently 100 µg of lysate was transferred to the wells in a black 96 well plate format. The plate was incubated for 1h at 37 ºC in the dark with reaction buffer and substrate. After which the fluorescence reading was made with a plate reader equipped with excitation at 400 nm and emission at 505 nm. Absorbance Signals in arbitrary units were presented after subtraction of background and were shown as the relative calpain activity.

### Western blot analysis

Western blot analysis was carried out according to a previously described standard protocol (52). Proteins from fracture femora were extracted by RIPA buffer combined with protease inhibitors and phosphatase inhibitors (Roche). The supernatants were obtained by centrifugation (12,000g, 10min, 4 °C). Protein concentration was determined utilizing the Pierce BCA Protein Assay kit. Equivalent amounts of protein in the specimen were separated based on their molecular weight by electrophoresis on 8% Bis-Tris gels and transferred to polyvinylidene difluoride (PVDF) membranes (Life Technologies). Membranes were immediately blocked with 5% nonfat dried milk diluted in TBST and incubated with primary antibodies (1:1,000, RFP from Rockland; *p-PI3K*, Cat: 17366s; *PI3K*, Cat: 4257s; *Akt*, Cat: 9272s; *p-Akt* Ser473, Cat: 9271s and *Notch1*, Cat: 4147 Val1744 D3B8, Rabbit mAb from Cell Signaling Technology; *CD31*, Cat: ab28364 from abcam) overnight at 4 °C. After 3 times rinses with TBST, membranes were incubated with horseradish peroxidase (HRP)-linked secondary antibodies. The protein bands were detected by ECL reagents (GE Healthcare) and the ChemiDoc XRS+ System (Bio-Rad). Duplicate experiments and analyses were performed at least 3 times.

### RT-PCR

Total RNA was extracted using a Tri-reagent protocol from femurs flushed to remove bone marrow followed by DNase I (Ambion) treatment. Reverse transcription (RT) was performed using a high capacity RNA-to-cDNA kit (Thermo Fisher, USA). The specificity of PCR was verified by performing reactions without RT and melt-curve analysis. Sequences of PCR primers were: Mouse *Piezo1* (forward), 5’-GCTTGCTAGAACTTCACG-3’, (reverse) 5’-GTACTCATGCGGGTTG-3’; Mouse *Hes1* (forward), 5’-CCTCTGAGCACAGAAAGTCA-3’, (reverse) 5’-GCCGGGAGCTATCTTTCTTA-3’; Mouse *Hey1* (forward), 5’-GTACCCAGTGCCTTTGAGAA-3’, (reverse) 5’-TTTCAGGTGATCCACAGTCA-3’. PCR products were electrophoresed on 1.5% agarose gels and sequenced to recognize their identity. PCR was performed on a CFX96 Real-Time PCR detection system (Bio-rad) using SYBR Green Premix Ex Taq II (Takara, Dalian, Liaoning, China). All the 2-ΔΔCT values were normalized using the reference gene β-actin.

### Isolation of liver endothelial cells

Murine liver tissues were kept in cold EBM-2 medium. Endothelial cells were isolated using CD31-conjugated microbeads (Miltenyi Biotec). Initially, the tissue was minced and resuspended in a disgestive solution consisting of 9 mL 0.1% collagenase II, 1 mL 2.5 U⋅mL ^−1^ dispase, 1 μM magnesium chloride and 1 µM calcium chloride in Hanks Buffer. The tissue-dissociation mix was incubated in a MACSMix Tube Rotator (Miltenyi Biotech) for 45 min at 37°C with continuous stirring. At the end of enzymatic dissociation, the sample was passed through 100 μm and 40 µm cell filters to remove undigested tissue, Cells were washed twice in magnetically activated cell sorting (MACS) buffer composed of PBS with 0.1% bovine serum albumin (BSA) and 2 mM EDTA at pH 7.2 followed by suspended in 20 mL red blood cell lysis buffer containing 0.206 g Tris base and 0.749 g NH_4_Cl in 100 mL PBS (pH 7.2) for 10 min, and then washed for a final time in MACS buffer. Next the pellet was incubated with 200 µL/1 × 10^7^ total cells of dead cell removal paramagnetic microbeads (Miltenyi Biotec) and incubated for 15 min at RT. After incubation, the cells were passed through a LS column prepared with 1× Binding Buffer (Miltenyi Biotec) in a magnetic field (MiniMACS Separator, Miltenyi Biotec). The elute was incubated with 30 µL FcR blocking reagent and 30 µL CD31-conjugated paramagnetic microbeads (Miltenyi Biotec) for 15 min at 4°C. After incubation, the solution was prepared with an MS column and MACS buffer. CD31-positive cells remained in the column, and CD31 negative cells passed through as eluents. CD31-positive cells were washed with warm EBM-2 medium, placed in one well of a fibronectin coated 6-well plate, and incubated in a 5% CO2 incubator. Medium was changed at 12 h, and then every 48 h until ready to be used.

### Intracellular calcium measurements

Liver endothelial cells were incubated for 1 h at 37°C with 1 µM Fura-2 AM (Thermo Fisher Scientific, USA) and 0.1% pluronic acid in Standard Bath Solution (SBS): 130 mM NaCl, 8 mM D-glucose, 5 mM KCl, 10 mM HEPES, 1.5 mM CaCl_2_ and 1.2 mM MgCl_2_, titrated to pH 7.4 with NaOH. Cells were washed three times in SBS. Measurements were made at RT (21 ± 2°C) and excited with 340 and 380 nm light, and emissions was collected at 510 nm. The change of intracellular Ca^2+^ concentration was expressed by the change of Fura-2 fluorescence ratio at two excitation wavelengths.

### Statistical analyses

All data were presented as the mean ± SEM, where n indicates the number of independent experiments. Unpaired, two-tailed Student’s t-tests were used in this study for comparing two groups. For some data sets, one-way ANOVA followed by Tukey’s multiple comparison test was used. No animals or samples were eliminated from analysis, and when applicable, animals were randomly assigned to treatment and control groups. In our experiments, *P<0.05* were regarded to be significant and indicated by ‘*’. The Origin software (8.5) was used for statistical analysis.

### Materials

Unless stated separately, all chemicals that used were purchased from Sigma-Aldrich. Fura-2-AM (molecular probe) was dissolved in DMSO at a concentration of 1 mM. Pluronic acid F127 was in DMSO of 10% w.v^−1^ and stored at RT. Yoda1 (Tocris) was prepared as a 10 mM stock solution.

## Conflict of interest

The authors have declared that no conflict of interest exists

## Acknowledgments

The study was supported by the National Natural Science Foundation of China (NSFC 81770453, 81774339, 81603641).

## Author contributions

PC, GZ, SJ ZW, QH conducted and analyzed the experiments. YN, BD, XP, SL, YH, RW and LZ helped in generation and management of mice breeding. GZ, LZ and SL performed the immunostaining experiments. GZ, SL, JY and LZ performed data analysis. HW provided intellectual input. JL conceived the idea for the project, generated research funds, led and coordinated the study. JL designed the study and wrote the paper with the help from PC. All authors reviewed the results, commented on the manuscript, and approved the final version of the manuscript.

